# Comparison of solitary and collective foraging strategies of *Caenorhabditis elegans* in patchy food distributions

**DOI:** 10.1101/744649

**Authors:** Siyu Serena Ding, Leah S. Muhle, André E. X. Brown, Linus J. Schumacher, Robert G. Endres

## Abstract

Collective foraging has been shown to benefit organisms in environments where food is patchily distributed, but whether this is true in the case where organisms do not rely on long-range communications to coordinate their collective behaviour has been understudied. To address this question, we use the tractable laboratory model organism *Caenorhabditis elegans*, where a social strain (*npr-1* mutant) and a solitary strain (N2) are available for direct comparison of foraging strategies. We first developed an on-lattice minimal model for comparing collective and solitary foraging strategies, finding that social agents benefit from feeding faster and more efficiently simply due to group formation. Our laboratory foraging experiments with *npr-1* and N2 worm populations, however, show an advantage for solitary N2 in all food distribution environments that we tested. We incorporated additional strain-specific behavioural parameters of *npr-1* and N2 worms into our model and computationally identified N2’s higher feeding rate to be the key factor underlying its advantage, without which it is possible to recapitulate the advantage of collective foraging in patchy environments. Our work highlights the theoretical advantage of collective foraging due to group formation alone without long-range interactions, and the valuable role of modelling to guide experiments.

## Introduction

Collective behaviour is displayed in many animal species including swarming insects, schooling fish, flocking birds, and troops of mammals (1–4). The effect of collective behaviour on foraging has been studied, with recent models and field experiments suggesting that collective search for food may improve food detection as well as food intake (5–8). For instance, computational models show that foraging in groups can provide an advantage for finding patchily (heterogeneously) distributed food, albeit using long-range interactions (9). While long-range interactions may apply to animals with good visual or acoustic senses (10, 11), this type of interaction may be less relevant for smaller mesoscopic animals with limited sensory modalities, including nematodes (roundworms), which are known to swarm (12) but whose collective foraging we know little about. Moreover, direct comparison between model predictions and experimental data is often limited by uncontrolled natural environments that the animals live in (13). Here we investigate the foraging strategies of *Caenorhabditis elegans*, a 1-mm long nematode with both collective and solitary foraging phenotypes. Experimental accessibility of *C. elegans* under controlled laboratory conditions further facilitates comparison with modelling outcomes.

*C. elegans* feed on bacteria that proliferate in rotten fruits and stems (14). The food resource in the worms’ natural environment fluctuates and is patchily distributed in space and time (15). Intriguingly, while *C. elegans* strains isolated from the wild exhibit varying degrees of collective feeding when cultured in the lab (16), the laboratory reference strain N2 feeds individually. This striking difference led us to hypothesise that the contrasting foraging strategies may confer advantages in the strains’ respective resource environments: Collective foraging may be beneficial for wild strains in their natural environments where food distribution is likely patchy, whereas solitary foraging may be better suited for the laboratory environment where food is much more homogeneous.

To test this hypothesis, we experimentally model solitary and collective behaviour with N2 (Figure 1a) and *npr-1* (Figure 1b) worms, respectively. The latter are N2 worms with a loss-of-function mutation *(ad609)* in the neuropeptide receptor gene *npr-1*, and are hyper-social with pronounced and persistent aggregate formation on food (16, 17). Thus N2 and *npr-1* worms represent opposite extremes of the *C. elegans* collective phenotype and provide a useful system for comparing solitary and collective foraging strategies in a genetic background that is identical except for the *npr-1* gene. Apart from regulating foraging, *npr-1* affects a suite of traits including the responses to O_2_, CO_2_, and pheromones (18–20). Past work examining the fitness consequences of these two strains either focus on the role of aggregation-independent behaviours such as dispersal and bordering in diverse food distribution environments (21), or the role of aggregation itself in relatively simple food environments (22). Therefore, the question remains how the solitary N2 and social *npr-1* strains perform in diverse and non-homogeneous resource environments, with contrasting collective foraging behaviours arising from group formation alone.

**Figure 1:**
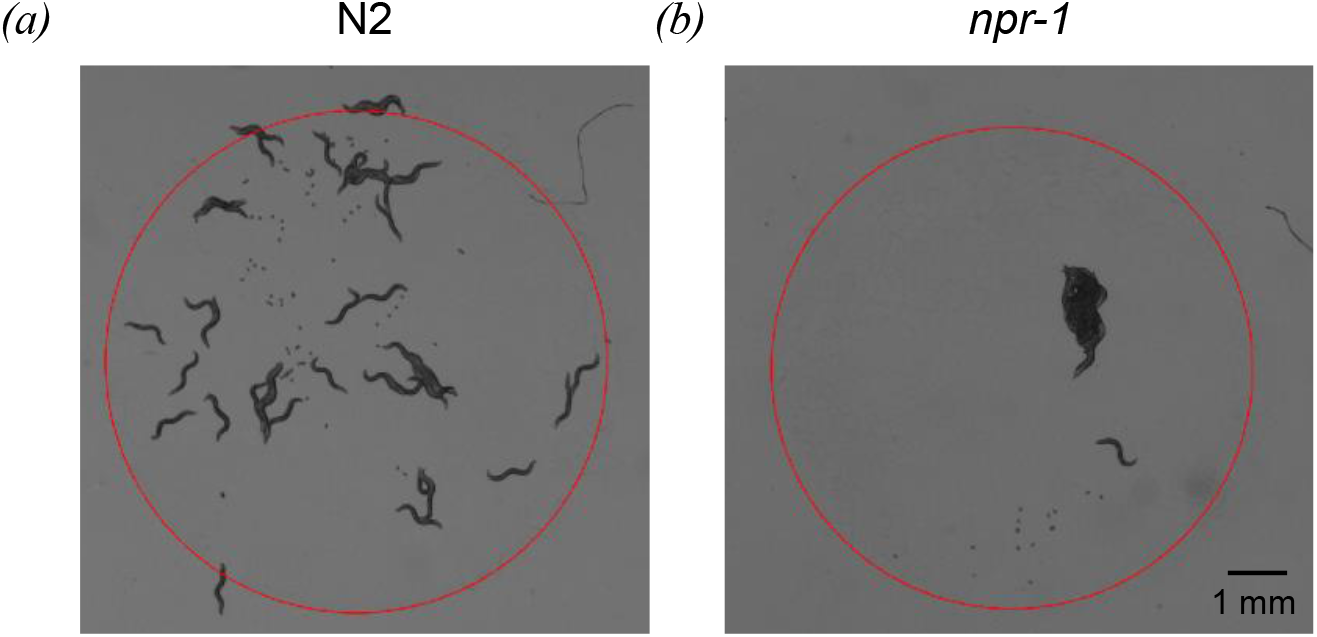
Snapshots of *C. elegans* on *E. coli* bacterial lawns from brightfield microscopy. *(a)* Solitary N2 worms on a bacterial lawn. *(b)* Hyper-social *npr-1(ad609)* worms on a bacterial lawn. Red circles indicate food boundaries, with food available only inside the circles.

To assess the effect of collective versus solitary foraging strategies in varying food environments, we developed a lattice-based foraging model for movement and feeding based on local interactions only. We first used a minimal model to investigate the sole effect of group formation on food, and then created a more realistic model that incorporates additional strain-specific behavioural parameters in order to facilitate direct comparison with the experimental data.

## Results

### Collective foraging is beneficial in patchy food distribution environments in the minimal model

To examine the exclusive effect of foraging in groups without considering any other behavioural differences, we first developed a minimal model where social and solitary agents are simulated to differ only in their ability to form groups on food. We use the terms “social” and “solitary” to refer to the individual propensity to aggregate, and “collective” and “solitary” to refer to the group-level foraging phenotypes. We refer to social individuals simply as those that aggregate and thus forage collectively, without any implication of complex social structure.

The basic agent behaviour in the minimal model is designed based on two observations from literature and from our experiments with both N2 and *npr-1* worms (Supplementary Movies S1-2). Firstly, worms move faster off food than on food, presumably to find new food (23). To implement this, at every time step both solitary and social agents move to one of eight lattice sites in the direct neighbourhood (to simulate slow movement) in the presence of food (Figure 2a, dark blue sites), or to one of sixteen sites in the remote neighbourhood (to simulate fast movement) in the absence of food (Figure 2a, light blue sites). In our model, an agent perceives food from the lattice site it is currently on and from the sites in its direct neighbourhood. The second experimental observation is that worms pump their pharynx and ingest bacteria whilst moving on food (24), which we simulate by having both types of agents consume one food unit per time step if they are on food.

**Figure 2:**
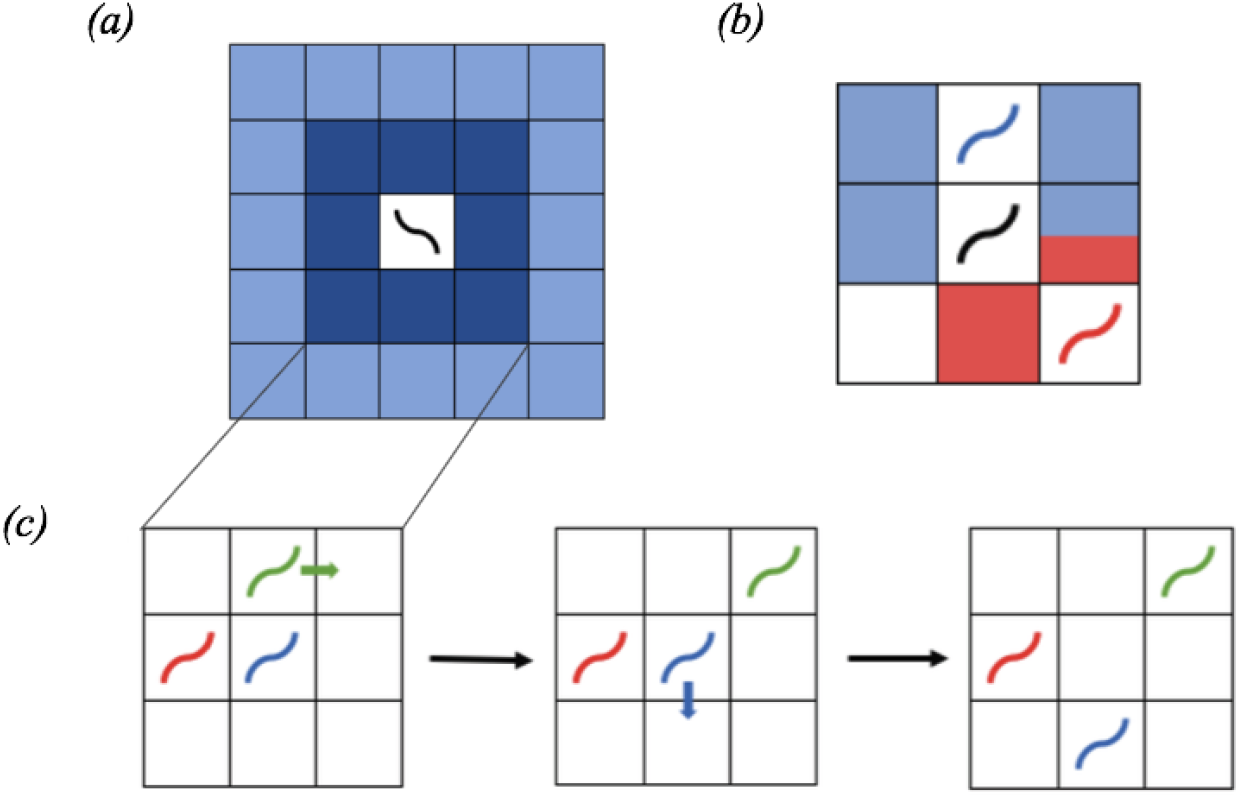
Schematics of neighbourhoods and computation of targeted steps. *(a)* Direct (dark blue) and remote (light blue) neighbourhoods of an agent (black worm) on a square lattice. *(b)* Possible motion updates of the black social agent performing a targeted step. Red sites show the direct neighbourhood shared by the red and the black agents, and blue sites show the direct neighbourhood shared by the blue and the black agents. Therefore, while performing a targeted step, the black agent is only allowed to move to one the five coloured sites (i.e. not the white sites), in order to perform a targeted step to the direct neighbourhood of an adjacent agent. *(c)* Consecutive execution of targeted steps in a group of three agents. The order in which motion updates are computed is chosen randomly for every time step. The green agent performs the first targeted step and moves to a square adjacent to the blue agent. Subsequently, the blue agent executes a targeted step and moves to a square next to the red agent which isolates the green agent from the group. This shows that a targeted step may also separate agents from their group.

The solitary and collective foraging strategies in the minimal model differ in the agents’ ability to form groups on food, and we implement this through the direction of agent movement. Social agents on food perform targeted steps towards neighbours (in order to form groups) if there are any in their direct neighbourhood (Figure 2b,c), otherwise all agents perform random steps (9) with step length determined by food availability. The minimal model simulations are thus constructed for examining exclusively the effect of neighbour attraction on foraging (see Materials and Methods for more details of the minimal model, and see Figure S1a for model flow chart). We chose to ignore long-range chemotaxis via food or pheromone signalling as our previous work suggests that these are not important for the aggregation phenotypes of the two worm strains (17).

We implement smoothly-varying, inhomogeneous food distributions with different degrees of food clustering controlled by a parameter *γ* in order to compare with previous work by Bhattacharya & Vicsek (9), based on which we constructed our minimal model but emphasising limited interaction range in our case. Each food unit is placed a distance *d* ≥ 1 away from an existing one with the probability *P*(*d*)~ *d*^−*γ*^ (see Materials and Methods). This parameterisation allows us to continuously vary between a uniformly random (*γ* = 0) food distribution and distributions with increasing patchiness as *γ* increases (Figure 3a).

**Figure 3:**
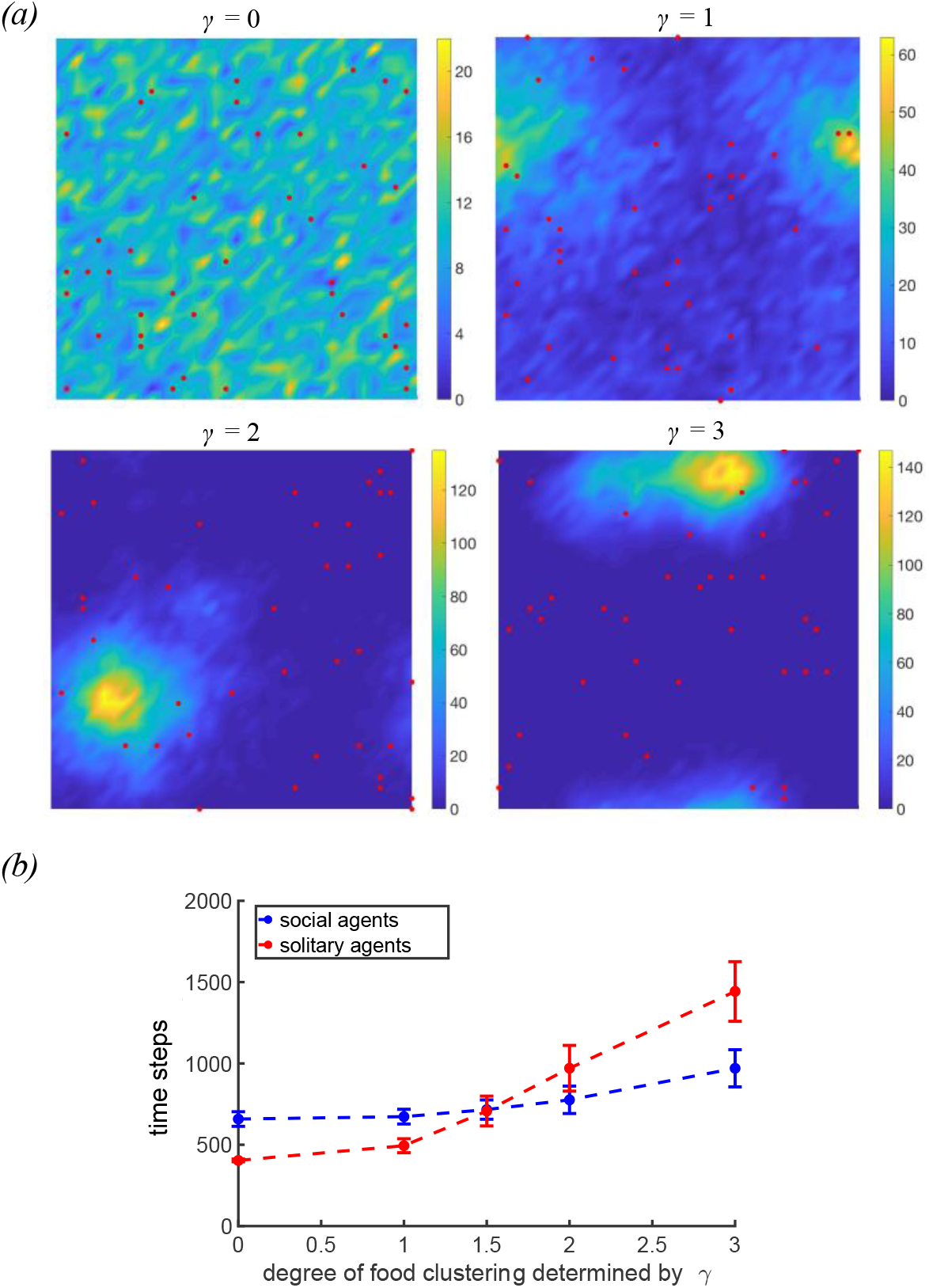
Minimal model simulations with smoothly-varying, inhomogeneous food distributions. *(a)* Food distributions for different *γ* values. Red dots show initial positions of the agents (distributed uniformly random), and the colour bars show the number of food units per lattice site. *(b)* Number of time steps taken by social and solitary agents to deplete 90% of the distributed food depending on the degree of food clustering, showing a crossover with social agents eating faster than solitary agents in patchy food environment (*γ*>1.5) and vice versa. Error bars show 1 SD.

In natural environments, *C. elegans* coexists with other bacterivores competing for the same food resources, so fast and efficient food depletion may enable a species to outperform its competitors (14, 25). Thus, we performed model simulations with populations of 40 agents and measured both time to 90% food depletion and foraging efficiency. In environments with uniform randomly distributed (*γ* = 0) or slightly patchy food (*γ* < 1.5), the solitary agents exhaust food faster than the social ones (Supplementary Movies S3-4); when food is more patchy (*γ* > 1.5), the reverse is true (Supplementary Movies S5-6). The crossover between the two foraging strategies can be found at approximately *γ* ≈ 1.5 (Figure 3b). Overall, these results support our initial hypothesis that a solitary foraging strategy is beneficial in environments with uniformly distributed food whereas collective foraging prevails in environments with patchy food. Interestingly, restricting food perception to the agent’s current lattice site diminishes the advantage of solitary agents in environments with uniformly random distributed or slightly patchy food (*γ* < 1.5) (Figure S2a).

The benefit of the collective foraging strategy can also be measured in terms of foraging efficiency, which is computed for individual agents by dividing the total number of food units it consumes by the total number of steps it takes; similar benefit-cost trade-offs had been considered by others in previous works (26, 27). In environments with uniformly random or slightly patchy food (*γ* < 1.5), solitary agents forage with a higher median efficiency than social ones, while the opposite is true in environments with patchy food distributions (*γ* > 1.5) (Figure S3a,b). However, the efficiencies of both social and solitary agents decrease as patchiness increases. Individual-level food consumption is less varied among solitary agents than among social ones in food environments with *γ* ≤ 1 (Figure S3c,d). With restricted food perception, however, the differences between agent types in individual efficiencies (Figure S2b,c) and individual food consumption (Figure S2d,e) disappear for *γ* ≤ 1.

These findings underline that collective foraging may be advantageous in environments with patchy food distribution due to both faster food consumption and higher foraging efficiency. The intuitive explanation for this is that in collective foraging the presence of other individuals may provide social information indicating the presence of food, like a queue forming at a conference buffet during lunch break. On a more abstract level, we can understand the advantage of collective foraging in patchy environments by considering the following: Initially, small aggregates may start to form anywhere in the environment. Aggregates at low food levels disperse more quickly as the food becomes depleted, whereas aggregates at high food levels persist longer, enabling aggregate growth as other agents join the group. Thus, social agents spend more time in regions with high food levels, leading to more successful foraging in patchy environments than the solitary agents who forage independently of other agents.

### Solitary N2 populations are more successful in laboratory foraging experiments

To test the predictions of the minimal model, we conducted population foraging experiments with social *npr-1(ad609)* mutants that feed in aggregates and solitary N2 worms that feed individually (Figure 1). We used food environments containing one, two, or four spots of *E. coli* OP50 bacteria (Figure 4a) to achieve increasing patchiness, because the smoothly-varying inhomogeneous distributions controlled by *γ* (Figure 3a) are difficult to produce experimentally. The total amount of bacteria remains the same across different experiments regardless of the spot number (i.e., 20 μL for one spot, 10 μL per spot for two spots and 5 μL per spot for four spots; see Materials and Methods). Note that a food “spot” is conventionally referred as a food “patch”, but here we use the term “spot” instead of “patch” to avoid confusion with the term “patchiness” (as opposed to uniformity), which in this context would refer to the presence of *multiple* spots (as opposed to a single spot). Each “spot” itself has a uniform distribution of food.

**Figure 4:**
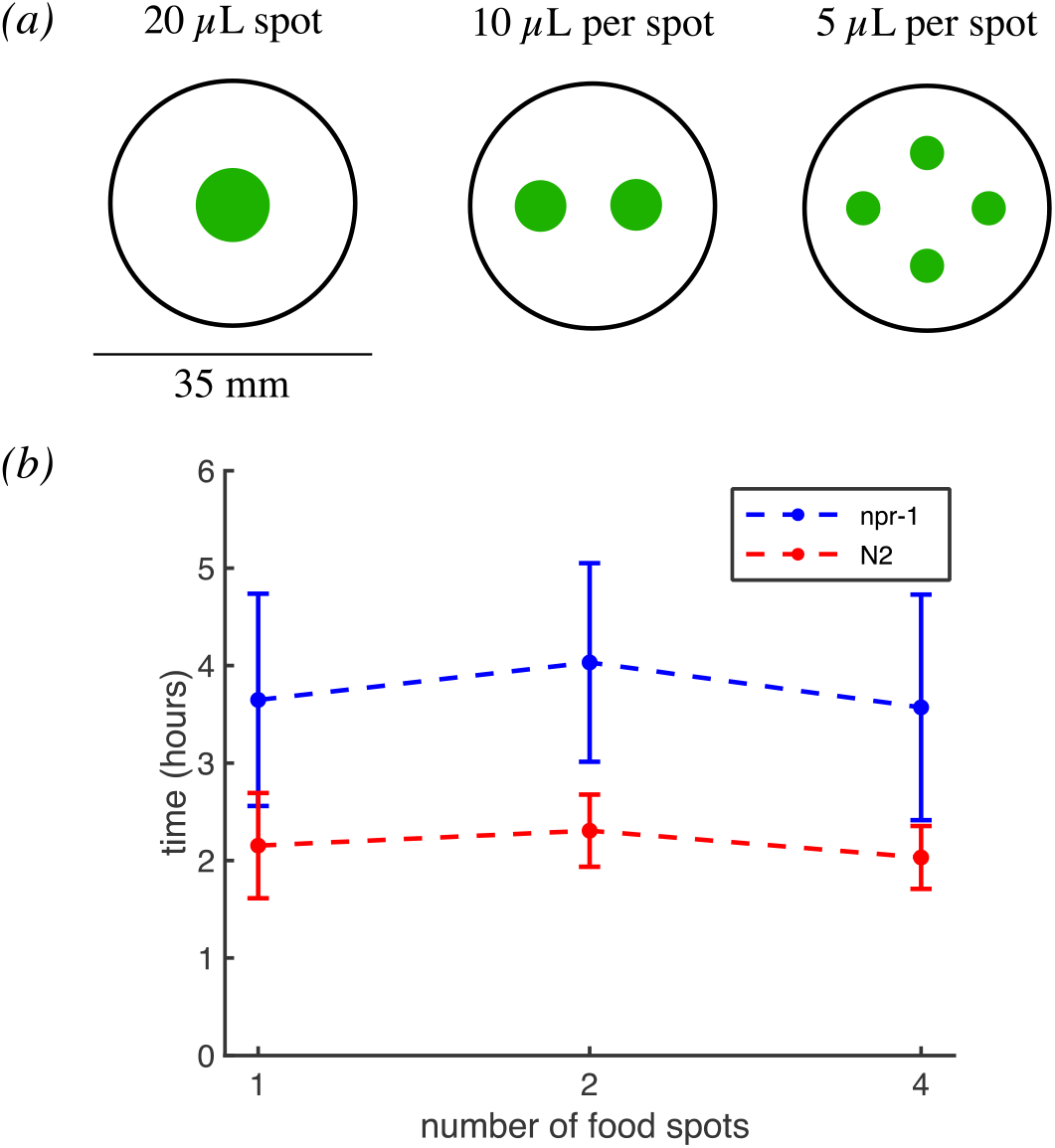
Experimental foraging assays with multi-spot food environments. *(a)* Schematics of food distributions in experiments. Shown are *E. coli* spots (green) on 35-mm Petri dishes with food spots arranged in one-, two-, and four-spot configurations. *(b)* Time for populations of 40 *npr-1* or N2 worms to exhaust food in the experiments. n = 5 independent replicates for each condition. Error bars show 1 SD.

We developed our experimental assay to circumvent the bordering and dispersal (i.e. leaving a food patch, instead of disbanding an aggregate) behaviours that Gloria-Soria & Azevedo (21) had previously focussed on, in order to assess the role of group formation on foraging success. We do so by using freshly seeded food spots to ensure that each spot has a uniform distribution without excessive bacterial growth in the border region. We also use a low level of peptone (0.013% w/v) in the media to minimise bacterial growth over the course of the experiment, which lasted up to seven hours. This foraging assay with thin, fresh bacterial lawns effectively eliminated bordering behaviour and led to very few food-leaving events. Food-leaving probability in our experiments are near zero (0.013 ± 0.013 (mean ± standard deviation) events per worm per hour for *npr-1* and 0.025 ± 0.025 events per worm per hour for N2, see Supplementary Methods), consistent with our previous report that worms are mostly on food under the same experimental conditions (17).

For experiments with either worm strain, each population consisted of 40 age-matched young adult worms. We measured the time taken to consume all the food in the environment. The end point of the assay is estimated from the detectable increase in worm speed once food becomes exhausted. This can be seen most clearly in Supplementary Movies S1-2, where the texture of the food patch changes from smooth to coarse upon consumption, and the drastic speed-up of the worms can be visually detected towards the end of both movies. Surprisingly, solitary N2 populations were faster at depleting the bacteria relative to *npr-1* populations independent of the number of bacteria spots (Figure 4b) (one spot: *npr-1* takes 70% longer than N2, two-sample t-test p = 0.01; two spots: *npr-1* takes 75% longer than N2, p < 0.01; four spots: *npr-1* takes 76% longer than N2, p = 0.01). Furthermore, time to food depletion barely varies amongst different food spot number configurations for both *npr-1* (one-way ANOVA p = 0.78) and N2 (p = 0.60) populations. Thus, the experimental results contradict the prediction of the minimal model, showing no advantage for collective feeding in patchy environments.

### Strain-specific model confirms experimental findings

In order to address the discrepancy between the minimal model predictions and our experimental findings, we created a more realistic, strain-specific version of the model, incorporating two more behaviours that differ between *npr-1* and N2 worms other than their tendency to form groups on food. Firstly, the speeds of *npr-1* and N2 worms differ depending on food availability. Both strains crawl at about the same speed in the absence of food; N2 worms slow down to roughly half this speed when on food, whereas *npr-1* worms only slow down significantly upon joining a group of worms on food (16). Secondly, *npr-1* worms have a feeding rate that is 62% that of N2, as calculated by us previously (28). These literature parameters are listed in Table 1 and adapted for our strain-specific simulations; model parameters are listed in Table 2. We do not use different food-leaving rates in our simulations because food-leaving is so rare in our experiments for both worm strains. Nevertheless, since others report much higher food-leaving rates under different experimental conditions (29, 30), our strain-specific model is constructed so that different food-leaving rates can easily be incorporated to test additional parameter combinations (see Supplementary Methods and Figure S1b for details). As in the minimal model, social agents (now called *npr-1* agents) on food join groups by performing targeted steps, whereas solitary agents (now called N2 agents) only perform random steps (see model flow chart in Figure S1b). In this strain-specific model, agents perceive food only from the lattice sites that they currently occupy.

**Table 1:** Literature values for *npr-1* and N2 behavioural parameters.

|  | reference | *npr-1* | N2 |
| --- | --- | --- | --- |
| **speed in the presence of food** | (16) | 183 $\mu\text{m/s}$ | 109 $\mu\text{m/s}$ |
| **speed in the absence of food** | (16) | 225 $\mu\text{m/s}$ | 232 $\mu\text{m/s}$ |
| **feeding rate** | (28) | 0.62 unit | 1 unit |

**Table 2:** Parameters used in modelling simulations.

|  | minimal model |  | strain-specific model |  |
| --- | --- | --- | --- | --- |
|  | social agents | solitary agents | *npr-1* agents | N2 agents |
| **Step length on food** | to direct neighbourhood | to direct neighbourhood | **in a group:** to direct neighbourhood<br>**alone:** to remote neighbourhood | to direct neighbourhood |
| **Step length off food** | to remote neighbourhood | to remote neighbourhood | to remote neighbourhood | to remote neighbourhood |
| **Feeding rate** | 1 food unit per time step | 1 food unit per time step | 0.4*0.62 food unit per time step* | 0.4 food unit per time step* |
| **Food-leaving probability** | Not used | Not used | 0 | 0 |

\* Feeding rates in the strain-specific model are scaled down to 0.4 for N2 and 62% of that for *npr-1*(28) to broadly match the experimental time to food depletion in Figure 5, based on a time step of $\Delta t = 10$ s.

We used multi-spot food distributions with one-, two-, or four-spot configurations in the strain-specific model (Figure 5a) to compare simulation outcomes with the experimental results. To assess foraging success in the strain-specific model, we calculated the time to 90% food depletion for both *npr-1* and N2 agent populations. N2 populations are faster at consuming the same amount of food than *npr-1* populations independent of the number of food spots (Figure 5b, Supplementary Movies S7-9), which confirms the experimental findings. We also analysed foraging efficiency of *npr-1* and N2 agents. These results show that N2 agents forage with a substantially higher efficiency than *npr-1* in all tested conditions, even though the range of individual efficiencies is larger for N2 (Figure S4a,c). However, *npr-1* agents have a higher median food intake than N2 in all environments, and fewer *npr-1* agents than N2 have an extremely low food intake (Figure S4b,d; Figure S5).

**Figure 5:**
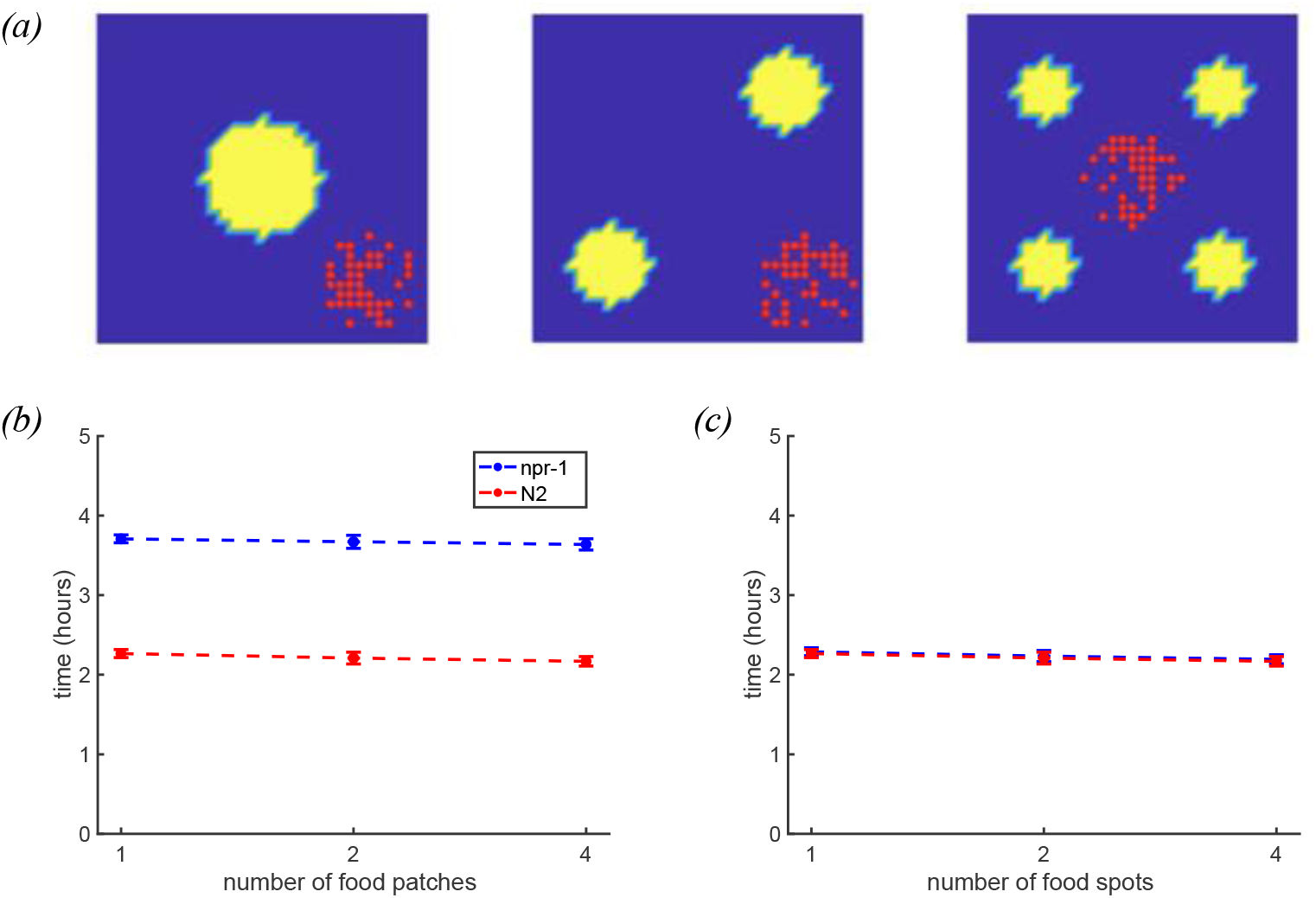
Strain-specific model simulations with multi-spot food environments. *(a)* Food distributions with one, two or four food spots. Red dots show agent configurations at the start of the simulations, clustered to mimic the experimental procedure of transferring worms together in a liquid droplet. Dark blue indicates no food and yellow indicates food. *(b)* Time for *npr-1* and N2 agents to deplete 90% of the distributed food units, shown for different numbers of food spots. Error bars show 1 SD. *(c)* Same as b), but with *npr-1* agent feeding rate set to the same value as N2. Simulation time is converted from time steps to real time in b) and c): As there is maximally a single agent per lattice site the lattice spacing is equal to the worm size (~ 1 mm). By noting that worm speed on food is approximately 100-200 μm/s and that it takes an agent one time step to cross the 1 mm lattice site, the timescale should be roughly ∆t ≈5-10 s. Eventually ∆t = 10 s is chosen to approximate the order of magnitude to broadly match the experimental data in Figure 4.

To ensure that the model outcome is not an artefact of using food environments consisting of distinct food spots, we repeated the strain-specific simulations with smoothly-varying inhomogeneous food distributions controlled by *γ*, as in the minimal model. We explored a broad range of *γ* values from 0 to 10, and confirmed that N2 agents still consume 90% of the food faster than *npr-1* agents for all tested food distributions (Figure 6a).

**Figure 6:**
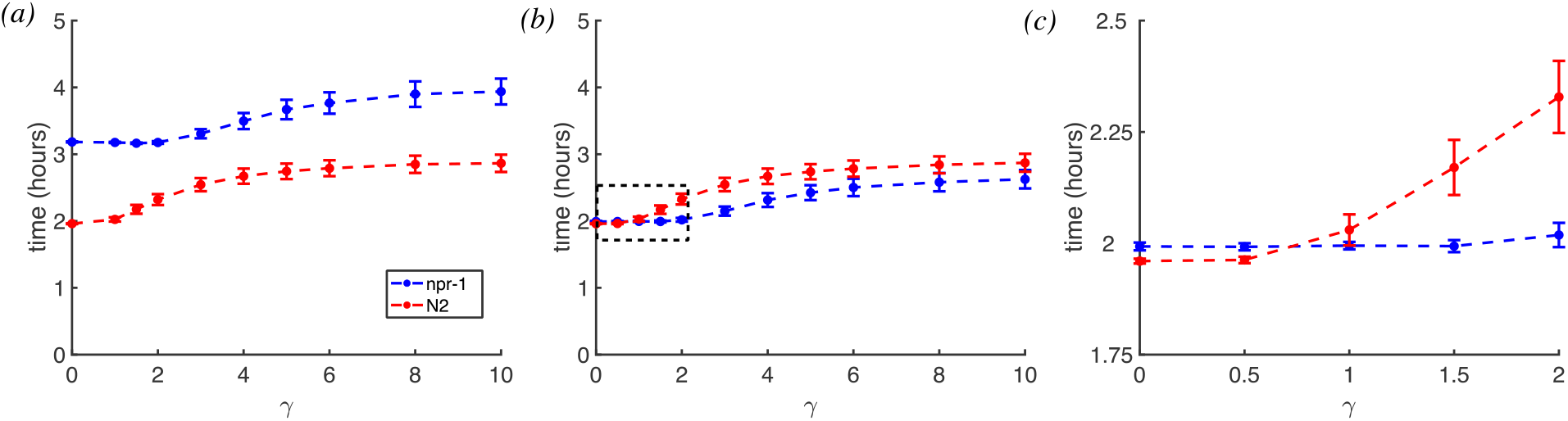
Strain-specific model simulations with smoothly-varying, inhomogeneous food distributions. *(a)* Time for *npr-1* and N2 agents to deplete 90% of the distributed food units, shown for different γ values. Error bars show 1 SD. Simulation time is converted from time steps to real time. *(b)* Same as a), but with *npr-1* agent feeding rate set to the same value as N2. The black dashed box is zoomed in and displayed in *(c)*. *(c)* Same as *(b)*, zoomed in to show a crossover of the agents’ foraging advantages between γ values of 0.5 and 1.

### Feeding rate is the key factor for N2’s foraging advantage

Now that we have a strain-specific model that matches our experimental data, we sought to determine which behavioural parameter underlies the difference between our minimal and strain-specific model outcomes. We repeated the strain-specific simulations with multi-spot food environments, but with equal feeding rates for *npr-1* and N2 agents (using the N2 value from Table 2). As a result, the difference between the strains in foraging time is completely abolished (Figure 5c). Furthermore, the distributions of individual efficiencies (Figure S4c,e) as well as of ingested food units (Figure S4d,f) for *npr-1* and N2 agents now resemble each other after setting the feeding rates equal. These results suggest that the higher feeding rate of N2 is the main reason for its foraging advantage in the strain-specific simulations.

Repeating these computational experiments using strain-specific simulations but with smoothly-varying inhomogeneous food distributions, we confirmed that setting *npr-1* and N2 feeding rates equal abolishes N2 agents’ foraging advantage for all but the lowest *γ* values (*γ* < 1), (Figure 6b-c). Interestingly, the crossover of foraging advantage that was previously seen in the minimal model (Figure 3b) now re-emerges (Figure 6c), with N2 agents having an advantage in environments with uniformly random or slightly patchy food (*γ* < 1) and *npr-1* agents performing better in environments with patchy food (*γ* ≥ 1). These results uncouple the dominating effect of N2’s higher feeding rate on the overall foraging success from other behavioural parameters, and demonstrate that an advantage of *npr-1* remains under patchy food conditions if not for its lower feeding rate.

## Discussion

Collective foraging may be beneficial for organisms in environments with patchy food distributions, but whether this also applies to organisms only relying on short-range communications to coordinate their collective behaviour has been unclear. We hypothesised that collective foraging in groups does confer such an advantage. To test this hypothesis we implemented lattice-based simulations, which are more computationally efficient than off-lattice agent-based models (17) or spatial Gillespie simulations (31), and have a long history in ecological modelling (26). Compared to Bhattacharya & Vicsek’s previous lattice-based simulations with long-range interactions over a distance many times the body size of an individual (9), we only allowed for short-range interactions in order to exclude the role of visual cues and long-range chemotaxis. In both cases, an advantage for collective foraging can be achieved, and our minimal model with only short-range interactions is more appropriate for cellular behaviour or that of nematodes such as *C. elegans*. Our approach is also different from other works which investigate optimal foraging in patchy environments based on the marginal value theorem (26, 27). Our minimal model supports our hypothesis that foraging in groups can be beneficial in environments with patchy food distributions, as social agents deplete food faster and more efficiently than solitary ones. Intuitively speaking, aggregation helps worms deplete a food patch before leaving it at the risk of not finding a new one. As food depletion leads to aggregate dispersal, groups of social worms will spend less time in low-food regions, and more time in high-food regions. Put differently, the simple presence of a worm may convey social information to other worms, such as indicating that food quality is sufficiently high (13, 32, 33). This type of swarm intelligence may be particularly valuable in the absence of sophisticated communication systems or long-range interactions.

In contrast to the minimal model, our more realistic strain-specific simulations show that the solitary N2 agents perform better than the social *npr-1* agents in all tested food distribution environments regardless of patchiness. Assuming fast food depletion as a fitness advantage, these results agree with a previous study reporting that the social strains are less fit in laboratory conditions (34). Moreover, a recent study shows that the observed fitness advantage of N2 over *npr-1* worms is in fact dissociable from their collective phenotypes (22). Indeed, we show that N2’s better foraging performance may be more attributable to its higher feeding rate than to its foraging strategy. Therefore even though our strain-specific model suggests that collective foraging is not a more efficient strategy, at least under our tested food distribution conditions, our minimal model and modified strain-specific model (with equal feeding rates and smoothly-varying inhomogeneous food) indicate that this remains a theoretical possibility.

Gloria-Soria & Azevedo have previously investigated how *npr-1* polymorphism in *C. elegans* can promote the co-existence of solitary and social foraging strategies in nature via resource partitioning (21). Central to their findings are the pronounced differences in bordering and dispersal (food-leaving) behaviours between the strains, both of which they show to be independent of aggregation. Here we developed an experimental assay to circumvent these two confounding behaviours, as well as computationally uncoupled the effect of feeding rate differences to reveal the underlying effect of foraging in groups on foraging success in diverse food environments. We show that foraging in groups may be beneficial in patchy food environments. Apart from foraging, aggregation into groups may also serve other ecologically-relevant functions such as protecting *C. elegans* from desiccation or UV radiation (35).

While using the model organism *C. elegans* enables us to conduct foraging experiments in controlled laboratory conditions, we were unable to experimentally demonstrate an advantage of collective foraging. Our simulation results suggest that two modifications may be necessary to achieve this. Firstly, we could compare the foraging performance of social *npr-1* worms to that of slow-feeding *eat* mutants in the solitary N2 genetic background (36), in order to remove the dominating effect of N2’s higher feeding rate. Secondly, using equal feeding rates in the strain-specific model, we only saw the re-emergence of collective foraging advantage in smoothly-varying inhomogeneous food distributions (Figure 6c) but not multi-spot environments (Figure 5c). This suggests that the multi-spot environments that we created experimentally and computationally were not patchy enough. We would thus need to discover experimentally accessible food distributions for which collective foraging has an advantageous in the context of our work, and testing various distributions with our strain-specific model can help explore such possibilities.

In summary, our simulations and experiments were designed to test whether collective foraging helps to consume patchily distributed food, which may be representative of resource distributions in the wild. While we conclude that it does in our minimal model, our experiments show that N2 populations outperform *npr-1* under all tested food distributions. By constructing a more realistic simulation incorporating strain-specific behavioural parameters, we were able to not only confirm experimental outcome but also computationally identify N2’s higher feeding rate as the main driver of its foraging advantage. Our simulations only considered spatial variation in the food distributions, but have not explored temporal fluctuations of the environment. The dynamics of environmental fluctuations have been shown to influence whether sensing or stochastic phenotype switching is favoured in growing populations (37). An alternative approach is to consider under what environmental conditions collective foraging strategies emerge by evolution (38). Thus the role of both fluctuating environments and evolution of foraging strategies are avenues for further theoretical work on the benefits of collective foraging strategies.

## Materials and Methods

### Basic simulation rules

The following rules apply to all simulations: We simulate (*n* = 40) agents on a square-lattice with *L*^2^ lattice sites (*L* = 35) using periodic boundary conditions (9). The direct neighbourhood of an agent is defined as the eight surrounding lattice sites, whereas the 16 lattice sites surrounding the direct neighbourhood are defined as the remote neighbourhood (Figure 2a). Each lattice site contains a certain number of food units depending on the underlying food distribution. Volume exclusion is enforced in all simulations so that every lattice site can only be occupied by a single agent. We use uniformly random initial positions of the agents for the minimal model (Figure 3a), and clustered initial position of the agents for the strain-specific model (Figure 5a) to better compare with experimental conditions. At every time step, each agent eats food if there are any at its current position, and attempts to move.

The order in which agents update their motion is randomly determined for every time step. All simulations were implemented with MATLAB R2018b. We ran the simulations 500 times for each condition, using different random initial distribution of agents for each simulation.

For every simulation the time taken to 90% food depletion is measured for the population, and the foraging efficiency and the total food uptake are measured for individual agents.

### Food distribution in simulations

Two different types of food distributions are used in the simulations. The first type (“smoothly-varying inhomogeneous”) has smoothly-varying inhomogeneous food distribution parameterised by *γ*, which controls the degree of clustering (Figure 3a) (9). For *γ* = 0 the food is distributed uniformly random on the lattice. For *γ* > 0, every new food unit is placed at a distance 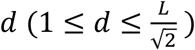 in a random direction to a random existing food unit. For *γ* > 0 the distance *d* is calculated as follows: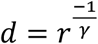, where *r* is a random number distributed uniformly between 0 and 1. If *d* is larger than 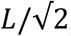, a uniform random value between 1 and 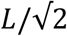 is chosen instead. The value of *d* is calculated independently for every food unit. To initialise simulations, one food unit is placed on a randomly chosen lattice site and then the remaining food units are distributed accordingly. There is a total of *L*^2^ · 10 food units in smoothly-varying inhomogeneous environments.

The second type of food distribution (“multi-spot”) consists of one, two, or four food spots distributed on the lattice, and food is distributed evenly between and within each spot (Figure 5a). The total food level is approximately *L*^2^ · 10 in multi-spot food environments, but varies slightly depending on the number of food spots, because each spot is made up of an integer number of lattice sites. To ensure consistent comparisons, we calculated the time to consuming *L*^2^ · 10 · 0.9 food units as time to depletion for every simulation.

### Minimal model simulations

Minimal model simulations are conducted with parameters listed in Table 2, and a flow chart is provided in Figure S1a. Different random initial food distributions are used for each simulation. Food is perceived from the lattice site that the agent currently occupies and from the eight sites in its direct neighbourhood. Social agents that are on food and has at least one other agent present in its direct neighbourhood perform a targeted step towards nearby neighbour(s) by moving randomly to one of the lattice sites located next to another agent in the direct neighbourhood (Figure 2b,c). Otherwise, all agents perform a random step to the direct neighbourhood if on food and to the remote neighbourhood if off food. The basics of random and targeted steps are also explained in the main text (Figure 2). An agent attempts movement into any unoccupied lattice site that fit the criteria, and if no such site is available, the agent remains at its current position. Agents eat one unit of food per time step if food is present. For the calculation of individual efficiencies, moving to the remote neighbourhood counts as two steps, moving to the direct neighbourhood counts as one step, and if the agent remains at its position then it counts as zero steps.

### Strain-specific model simulations

The parameters for strain-specific simulations including movement speeds and feeding rates are given in Table 2, and a flow chart is provided in Figure S1b. The initial food distribution for the strain-specific simulations is identical for each simulation (uniform spots), to mimic experimental conditions. In these simulations, an agent perceives food only from the lattice site that it currently occupies. Agents perform targeted and random steps in the same way as in the minimal models. The strain-specific model also incorporates food-leaving probability ρ (see Supplementary Methods), which is set to zero for our simulation results here. Foraging efficiencies for strain-specific simulations are calculated in the same way as in the minimal model simulations.

### Experimental procedure to validate the strain-specific simulations

The experimental procedures used here are identical to the “Bright field standard swarming imaging” method that we previously published (17). A step-by-step protocol is available at http://dx.doi.org/10.17504/protocols.io.vyhe7t6. Briefly, 35-mm imaging plates containing low peptone (0.013% w/v) NGM agar were seeded with 20 μL of diluted *E. coli* OP50 bacteria (OD_600_ = 0.75) shortly before imaging, with the 20 μL equally divided between the required number of food spots to produce different patchiness conditions (i.e., four spots of 5 μL each, two spots of 10 μL each, or one spot with 20 μL). Only freshly seeded (< 2 hours) plates were used for imaging to ensure food uniformity within each food spot, as long incubation would lead to a thicker border region due to bacteria growth. Forty age-synchronised young adult worms are washed and transferred onto the imaging plate in a liquid drop away from the bacterial spots, and imaging commences immediately. Time-lapse images were recorded at 25 fps for 7 hours at 20° C with Gecko software (v2.0.3.1) and a custom-built six-camera rig equipped with Dalsa Genie cameras (G2-GM10-T2041). Five replicates of the experiments are available for each combination of worm strain and food distribution condition, and the data are available at https://doi.org/10.5281/zenodo.3625159.

### Estimating the time to food depletion from experimental data

Time to food depletion was defined as the time difference between foraging start and complete food exhaustion, and these points were identified by visual assessment of recorded experiments. As worms were transferred to the imaging plates in a liquid drop that prevents escape, we defined foraging start time as the moment that the liquid drop is completely absorbed into the media allowing all worms to crawl out. As for the end point of food depletion, we identified drastic increases in overall worm speeds in our recordings as a proxy, because worms visibly speed up when food becomes exhausted. Such speed increases can occur more than once when multiple food spots exist as not all spots become simultaneously exhausted; we used the final instance to identify the point of total food depletion from all food spots.

## Acknowledgements

We would like to thank Camille Straboni from the Brown lab for providing additional brightfield movies. Some strains were provided by the CGC, which is funded by NIH Office of Research Infrastructure Programs (P40 OD010440).

## Supplementary Material

Supplementary Materials accompany this publication, including 13 Supplementary Movies, 5 Supplementary Figures, 1 Supplementary File, 3 Supplementary Methods, and 3 Supplementary References.

## Data accessibility

The code for the model simulations, the simulation results, and the code for the analysis of these results can be found at https://github.com/lsmuhle/CollectiveFeeding. Experimental recordings are available at https://doi.org/10.5281/zenodo.3625159 as part of the Open Worm Movement Database. The source file for the experimental analysis is listed as Supplementary File S1.

## Author contributions

SSD performed the experiments and conducted the analysis of the experimental data. LSM, LJS and RGE designed the models. LSM implemented the models and conducted the analysis of the simulations. LJS, RGE, and AEXB supervised the project. The manuscript was written by SSD, LJS, and LSM. SSD and LJS led critical revision of the manuscript.

## References

1. Partridge BL. The structure and function of fish schools. Sci Am. 1982;246(6):114–23.

2. Emlen JT, Jr. Flocking behavior in birds. The Auk. 1952;69:160–70.

3. Simpson SJ, Sword GA. Locusts. Current Biology. 2008;18(9).

4. Strandburg-Peshkin A, Farine DR, Couzin ID, Crofoot MC. Shared decision-making drives collective movement in wild baboons. Science. 2015.

5. Snijders L, Kurvers RHJM, Krause S, Ramnarine IW, Krause J. Individual- and population-level drivers of consistent foraging success across environments. bioRxiv. 2018.

6. Li L, Peng H, Kurths J, Yang Y, Schellnhuber HJ. Chaos-order transition in foraging behavior of ants. Proceedings of the National Academy of Sciences of the United States of America. 2014;111(23):8392–7.

7. Torney CJ, Berdahl A, Couzin ID. Signalling and the Evolution of Cooperative Foraging in Dynamic Environments. Plos Comput Biol. 2011;7(9).

8. Cvikel N, Berg KE, Levin E, Hurme E, Borissov I, Boonman A, et al. Bats Aggregate to Improve Prey Search but Might Be Impaired when Their Density Becomes Too High. Current Biology. 2015;25(2):206–11.

9. Bhattacharya K, Vicsek T. Collective foraging in heterogeneous landscapes. J R Soc Interface. 2014;11(100).

10. Beauchamp G. Exploring the role of vision in social foraging: what happens to group size, vigilance, spacing, aggression and habitat use in birds and mammals that forage at night? Biol Rev Camb Philos Soc. 2007;82(3):511–25.

11. Gager Y, Gimenez O, O’Mara MT, Dechmann DK. Group size, survival and surprisingly short lifespan in socially foraging bats. BMC Ecol. 2016;16:2.

12. McBride JM, Hollis JP. Phenomenon of Swarming in Nematodes. Nature. 1966;211(5048):545–6.

13. Berdahl AM, Kao AB, Flack A, Westley PAH, Codling EA, Couzin ID, et al. Collective animal navigation and migratory culture: from theoretical models to empirical evidence. Philos T R Soc B. 2018;373(1746).

14. Felix MA, Braendle C. The natural history of Caenorhabditis elegans. Curr Biol. 2010;20(22):R965–9.

15. Frezal L, Felix MA. C. elegans outside the Petri dish. eLife. 2015;4.

16. de Bono M, Bargmann CI. Natural variation in a neuropeptide Y receptor homolog modifies social behavior and food response in C. elegans. Cell. 1998;94(5):679–89.

17. Ding SS, Schumacher LJ, Javer AE, Endres RG, Brown AE. Shared behavioral mechanisms underlie C. elegans aggregation and swarming. eLife. 2019;8.

18. Gray JM, Karow DS, Lu H, Chang AJ, Chang JS, Ellis RE, et al. Oxygen sensation and social feeding mediated by a C. elegans guanylate cyclase homologue. Nature. 2004;430(6997):317–22.

19. Macosko EZ, Pokala N, Feinberg EH, Chalasani SH, Butcher RA, Clardy J, et al. A hub-and-spoke circuit drives pheromone attraction and social behaviour in C. elegans. Nature. 2009;458(7242):1171–5.

20. Rogers C, Persson A, Cheung B, de Bono M. Behavioral motifs and neural pathways coordinating O2 responses and aggregation in C. elegans. Curr Biol. 2006;16(7):649–59.

21. Gloria-Soria A, Azevedo RB. npr-1 Regulates foraging and dispersal strategies in Caenorhabditis elegans. Curr Biol. 2008;18(21):1694–9.

22. Zhao Y, Long L, Xu W, Campbell RF, Large EE, Greene JS, et al. Changes to social feeding behaviors are not sufficient for fitness gains of the Caenorhabditis elegans N2 reference strain. eLife. 2018;7.

23. Gray JM, Hill JJ, Bargmann CI. A circuit for navigation in Caenorhabditis elegans. Proceedings of the National Academy of Sciences of the United States of America. 2005;102(9):3184–91.

24. Scholz M, Dinner AR, Levine E, Biron D. Stochastic feeding dynamics arise from the need for information and energy. Proceedings of the National Academy of Sciences of the United States of America. 2017;114(35):9261–6.

25. Riddle DL, Blumenthal T, Meyer BJ, Priess JR. Introduction to C. elegans. In: nd, Riddle DL, Blumenthal T, Meyer BJ, Priess JR, editors. C elegans II. Cold Spring Harbor (NY)1997.

26. Charnov EL. Optimal foraging, the marginal value theorem. Theor Popul Biol. 1976;9(2):129–36.

27. Wajnberg E, Fauvergue X, Pons O. Patch leaving decision rules and the Marginal Value Theorem: an experimental analysis and a simulation model. Behavioral Ecology. 2000;11(6):577–86.

28. Ding SS, Romenskyy M, Sarkisyan KS, Brown AEX. Measuring Caenorhabditis elegans Spatial Foraging and Food Intake Using Bioluminescent Bacteria. Genetics. 2020.

29. Milward K, Busch KE, Murphy RJ, de Bono M, Olofsson B. Neuronal and molecular substrates for optimal foraging in Caenorhabditis elegans. Proceedings of the National Academy of Sciences of the United States of America. 2011;108(51):20672–7.

30. Shtonda BB, Avery L. Dietary choice behavior in Caenorhabditis elegans. J Exp Biol. 2006;209(Pt 1):89–102.

31. Abel JH, Drawert B, Hellander A, Petzold LR. GillesPy: A Python Package for Stochastic Model Building and Simulation. IEEE Life Sci Lett. 2016;2(3):35–8.

32. Krause J, Ruxton GD, Krause S. Swarm intelligence in animals and humans. Trends Ecol Evol. 2010;25(1):28–34.

33. Hein AM, Rosenthal SB, Hagstrom GI, Berdahl A, Torney CJ, Couzin ID. The evolution of distributed sensing and collective computation in animal populations. eLife. 2015;4:e10955.

34. Andersen EC, Bloom JS, Gerke JP, Kruglyak L. A variant in the neuropeptide receptor npr-1 is a major determinant of Caenorhabditis elegans growth and physiology. PLoS Genet. 2014;10(2):e1004156.

35. Busch KE, Olofsson B. Should I stay or should I go? Worm. 2012;1(3):182–6.

36. Avery L. The Genetics of Feeding in Caenorhabditis elegans. Genetics. 1993;133:897–917.

37. Kussell E, Leibler S. Phenotypic diversity, population growth, and information in fluctuating environments. Science. 2005;309(5743):2075–8.

38. Witkowski O, Ikegami T. Emergence of Swarming Behavior: Foraging Agents Evolve Collective Motion Based on Signaling. Plos One. 2016;11(4).

