## Supplementary Information for "Comparison of solitary and collective foraging strategies of *Caenorhabditis elegans* in patchy food distributions"

Including:

13 Supplementary Movies,

5 Supplementary Figures,

1 Supplementary File,

3 Supplementary Methods, and

3 Supplementary References

### Supplementary Movies

Supplementary movies can be viewed at https://tinyurl.com/y5bvolq3.

**Movie S1: Sample brightfield recording of N2 worms feeding on a bacterial lawn.**

**Movie S2: Sample brightfield recording of *npr-1* worms feeding on a bacterial lawn.**

**Movie S3: Minimal model simulation of social (left) and solitary (right) agents in an environment with uniformly random distributed food ($\gamma = 0$).** Red dots represent the forty agents, dark blue indicates no food and lighter blue/green/yellow show increased food levels.

**Movie S4: Minimal model simulation of social (left) and solitary (right) agents in an environment with slightly patchy food ($\gamma = 1$).** Red dots represent the forty agents, dark blue indicates no food and lighter blue/green/yellow show increased food levels.

**Movie S5: Minimal model simulation of social (left) and solitary (right) agents in an environment with strongly patchy food ($\gamma = 2$).** Red dots represent the forty agents, dark blue indicates no food and lighter blue/green/yellow show increased food levels.

**Movie S6: Minimal model simulation of social (left) and solitary (right) agents in an environment with very strongly patchy food ($\gamma = 3$).** Red dots represent the forty agents, dark blue indicates no food and lighter blue/green/yellow show increased food levels.

**Movie S7: Strain-specific model simulation of *npr-1* (left) and N2 (right) agents in an environment with one food spot.** Red dots represent the forty agents, with dark blue indicating no food and yellow indicating food.

**Movie S8: Strain-specific model simulation of *npr-1* (left) and N2 (right) agents in an environment with two food spots.** Red dots represent the forty agents, with dark blue indicating no food and yellow indicating food.

**Movie S9: Strain-specific model simulation of *npr-1* (left) and N2 (right) agents in an environment with four food spots.** Red dots represent the forty agents, with dark blue indicating no food and yellow indicating food.

**Movie S10: Strain-specific model simulation of *npr-1* (left) and N2 (right) agents in an environment with uniformly random distributed food ($\gamma = 0$).** Red dots represent the forty agents, dark blue indicates no food and lighter blue/green/yellow show increased food levels.

**Movie S11: Strain-specific model simulation of *npr-1* (left) and N2 (right) agents in an environment with slightly patchy food ($\gamma = 1$).** Red dots represent the forty agents, dark blue indicates no food and lighter blue/green/yellow show increased food levels.

**Movie S12: Strain-specific model simulation of *npr-1* (left) and N2 (right) agents in an environment with strongly patchy food ($\gamma = 2$).** Red dots represent the forty agents, dark blue indicates no food and lighter blue/green/yellow show increased food levels.

**Movie S13: Strain-specific model simulation of *npr-1* (left) and N2 (right) agents in an environment with very strongly patchy food ($\gamma = 3$).** Red dots represent the forty agents, dark blue indicates no food and lighter blue/green/yellow show increased food levels.

### Supplementary Figures

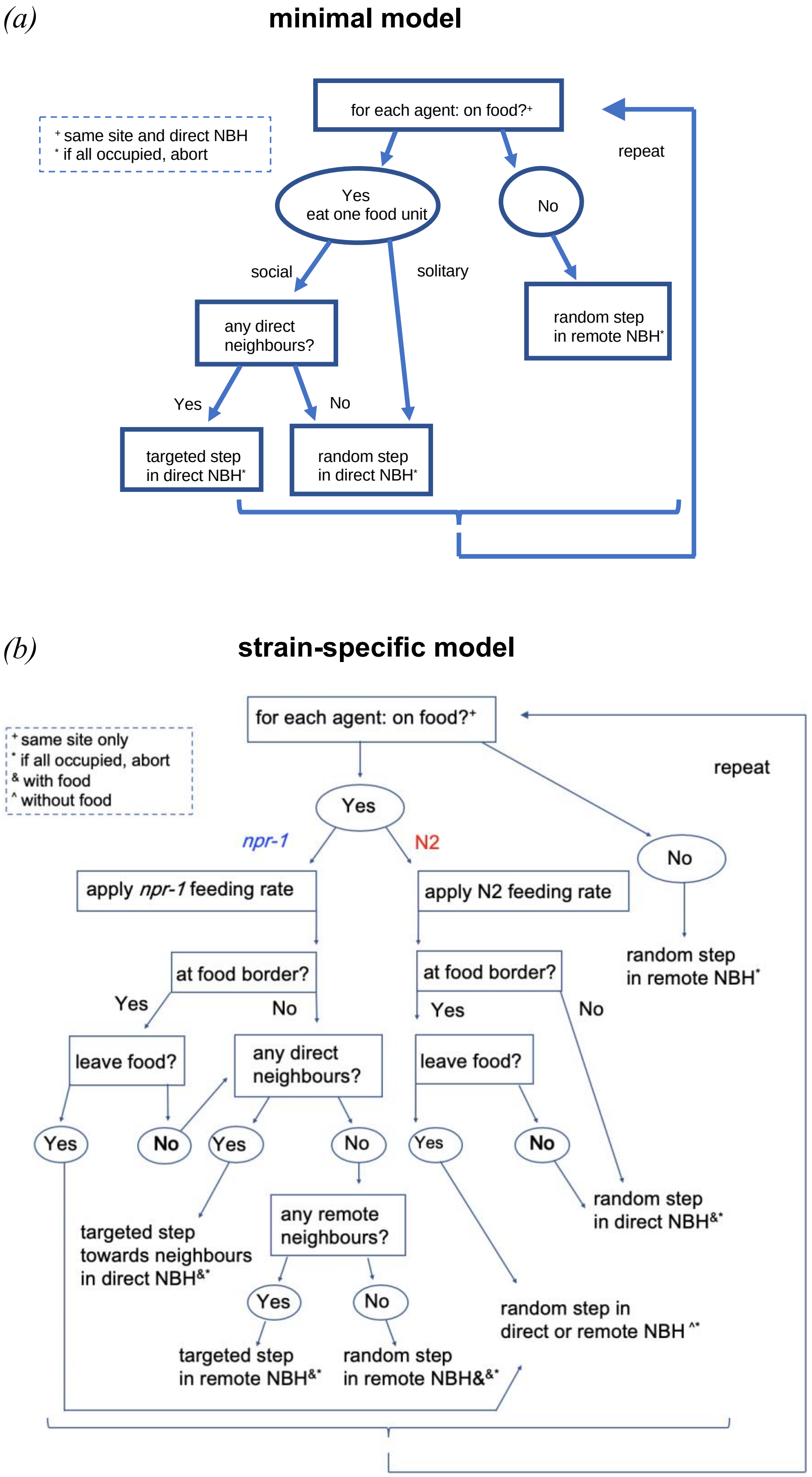


**Figure S1: Flow charts of models.** Parameters of (*a*) the minimal model and (*b*) the strain-specific model are based on Table 1 and listed in Table 2. In (*b*), bold and regular fonts correspond to high and low probabilities, respectively. NBH = neighborhood.

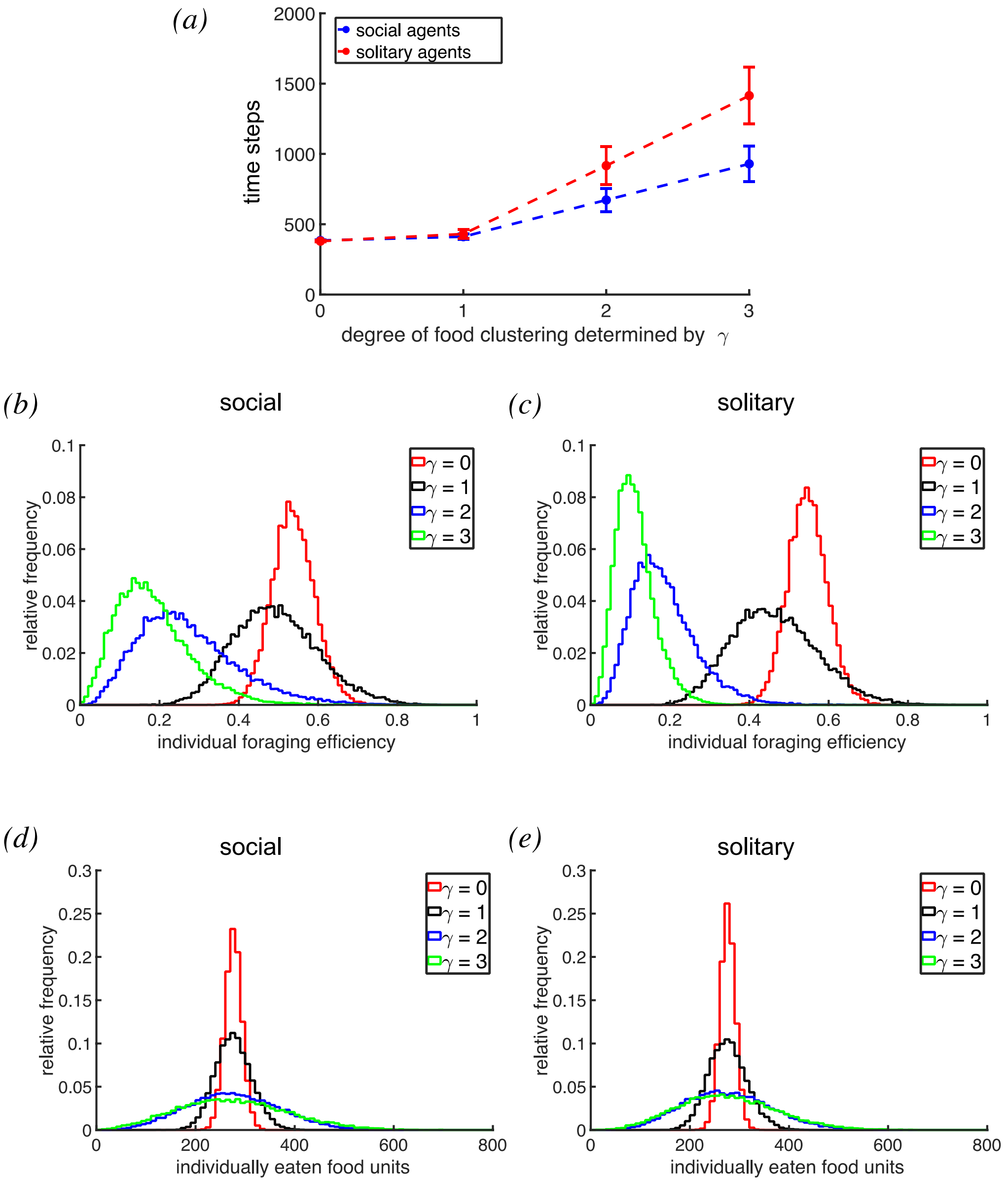


**Figure S2: Characterisation of social and solitary agents in the minimal model with restricted food perception.** Restricted food perception here means agents perceive food only on their current lattice site, but not in their direct neighbourhood. *(a)* Number of time steps taken by social and solitary agents to deplete 90% of the distributed food. Error bars show 1 SD. Distributions of *(b)* individual efficiencies of social agents, *(c)* individual efficiencies of solitary agents, *(d)* ingested food units of social agents and *(e)* ingested food units of solitary agents.

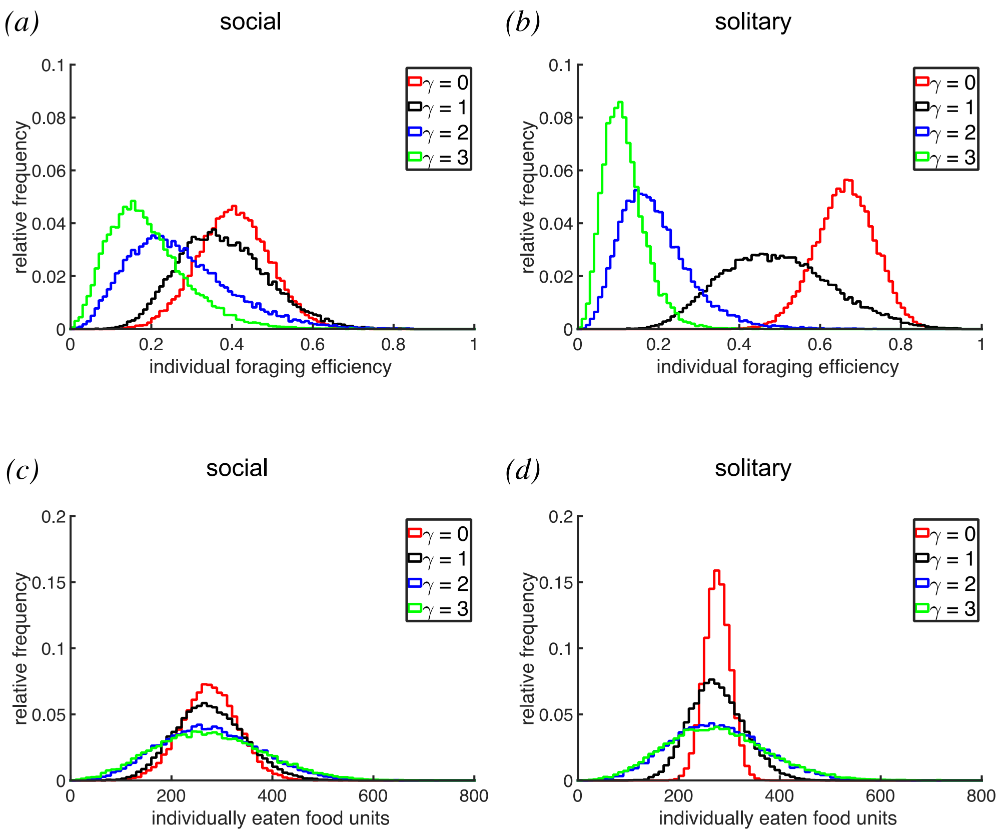


**Figure S3: Distributions of individual foraging efficiencies and total food uptake.** Shown are social and solitary agents in the minimal model simulations with dependence on the degree of food clustering (determined by parameter $\gamma$). Distributions of *(a)* individual efficiencies of social agents, *(b)* individual efficiencies of solitary agents, *(c)* ingested food units of social agents and *(d)* ingested food units of solitary agents.

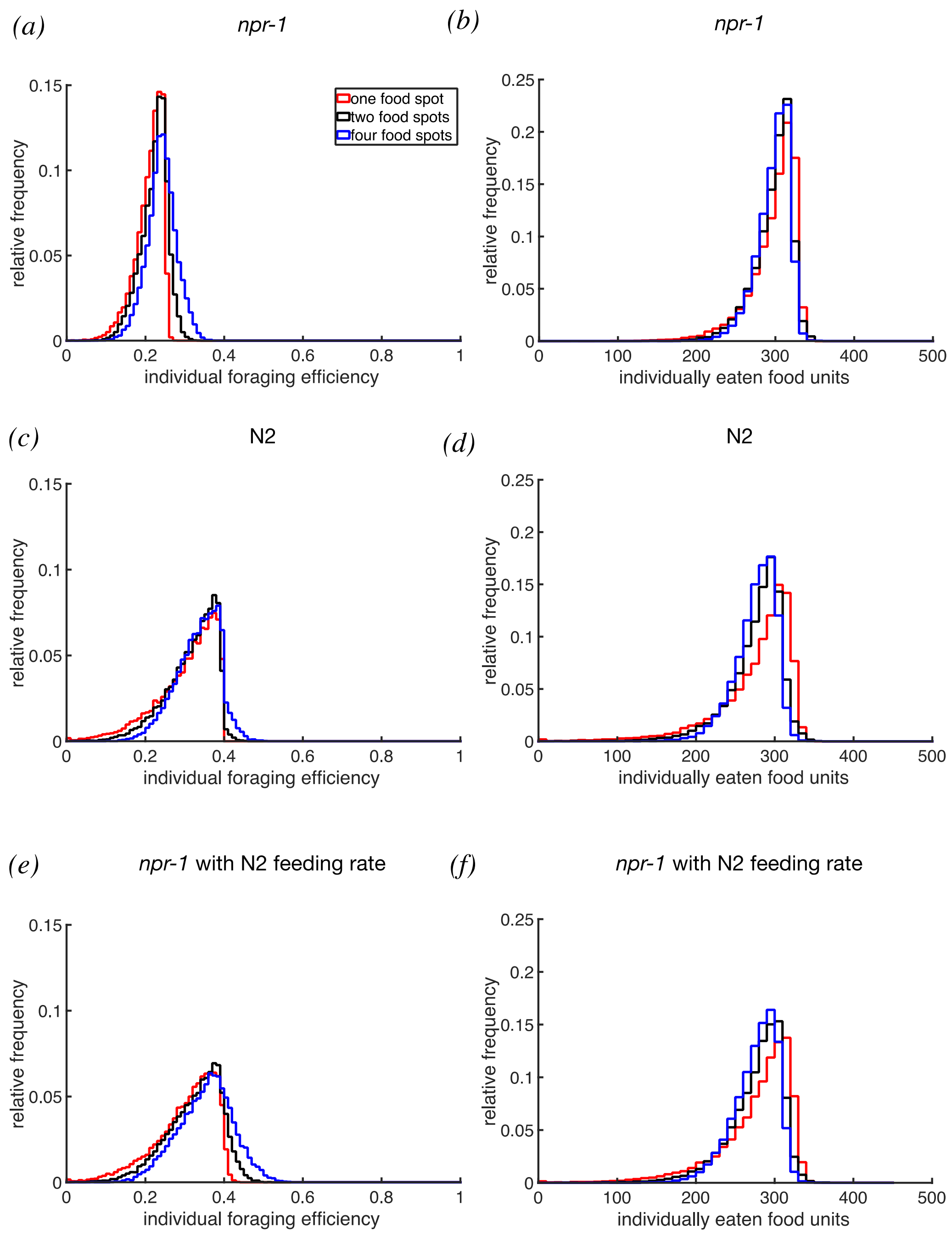


**Figure S4: Distributions of individual foraging efficiencies and total food uptake.** Shown are *npr-1* and N2 agents in environments with one, two or four food spots in the strain-specific simulations. Distributions of *(a)* individual efficiencies of *npr-1* agents, *(b)* individual efficiencies of N2 agents, *(c)* ingested food units of *npr-1* agents, *(d)* ingested food units of N2 agents, *(e)* individual efficiencies of *npr-1* agents with the same feeding rate as N2 agents (0.4 units per time step) and *(f)* ingested food units of *npr-1* agents with the same feeding rate as N2 agents (0.4 units per time step).

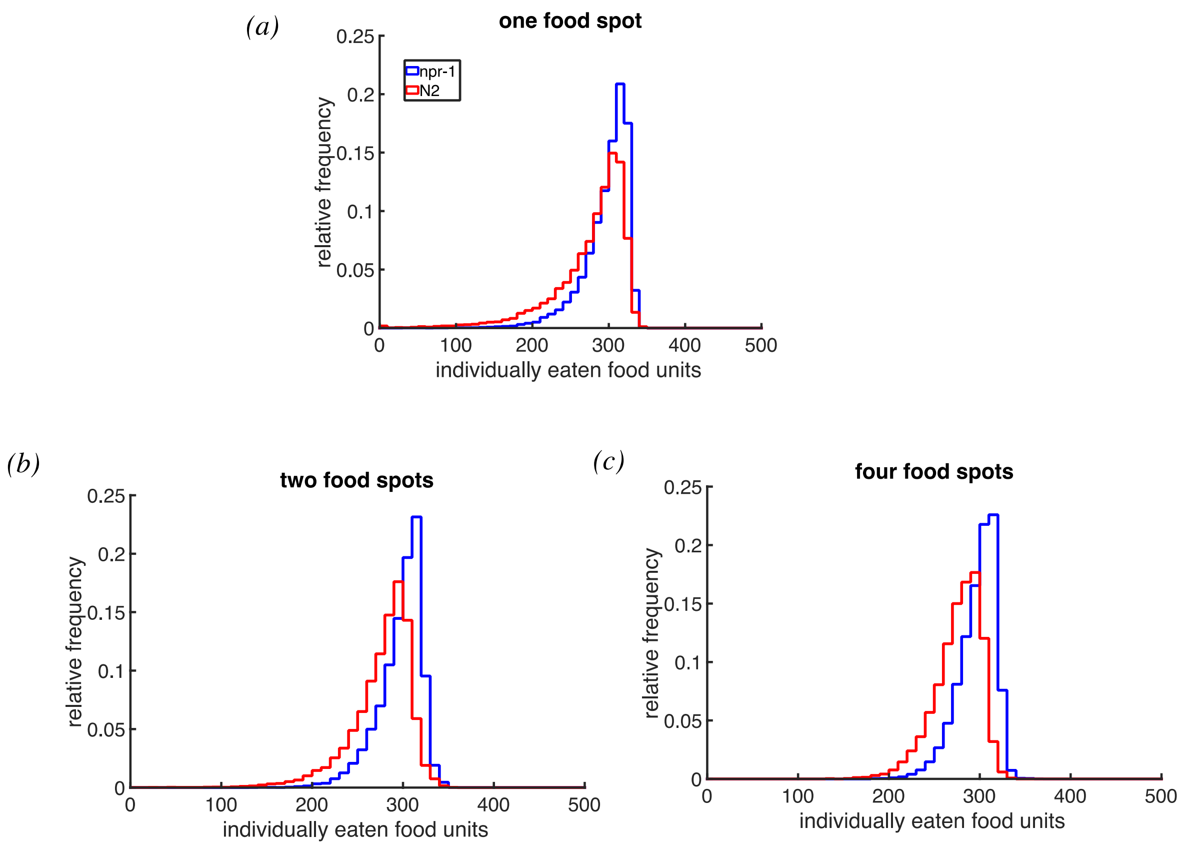


**Figure S5: Distributions of total food uptake of *npr-1* and N2 agents in direct comparison.** Shown are the distributions of number of ingested food units per agent in an environment with *(a)* one food spot, *(b)* two food spots and *(c)* four food spots in the strain-specific simulation.

### Supplementary File

**File S1:** Source file for the analysis of foraging experiments with N2 and *npr-1* worm populations.

### Supplementary Methods

#### *Details on implementing food-leaving for the strain-specific model*

To implement the food-leaving probability ρ, the food border needs to be defined. An on-food N2 agent is considered to be at the food border if there is at least one lattice site without food in the direct neighbourhood. An on-food *npr-1* agent is considered to be at the food border in one of the following three scenarios: a) if it is in a group (having at least one neighbour in the direct neighbourhood), and if there is at least one lattice site without food in the direct neighbourhood; b) it has no direct neighbours, but has at least one neighbour and at least one lattice site without food in the remote neighbourhood; c) it has no neighbours in either the direct or the remote neighbourhood, and has at least one lattice site without food in the remote neighbourhood. Thus, *npr-1* agents have an increased food perception in this context, because when they move at higher speeds they are aware of whether moving to a site with or without food. Similarly, *npr-1* agents have an increased perception of other agents on food. Agents without direct neighbours look for neighbours in the remote neighbourhood and, if there are any, perform a targeted step to the remote neighbourhood. *npr-1* agents in a group on food perform targeted steps to the direct neighbourhood (see below for details on motion updates).

#### *Details on motion updates for the strain-specific model*

A flow chart (Supplementary Figure S1b) shows the schematics of the following motion updates.

For *npr-1*, an agent that is located at the food border decides according to the food-leaving probability ρ whether to remain on food or leave it. For this, a random number $r$ uniformly distributed between 0 and 1 is drawn. If $r \leq \rho$, the agent moves to a field without food, whereas it moves to a field with food if $r > \rho$. The computation of the random step in the absence of food is the same as in the minimal model simulations. If the agent is located on food, not at the food border and has no direct or remote neighbours, it performs a random step to the remote neighbourhood. If it is on food, not at the food border and only has neighbours in the remote neighbourhood, it performs a targeted step to the remote neighbourhood. However, if the agent is on food, not at the food border, but has direct neighbours, it performs a targeted step to the direct neighbourhood. In the case that an agent is located on food and at the food border, the computations differ. If it leaves the food, it can move to all lattice sites without food in the direct or remote neighbourhood depending on its speed. If no lattice site without food is unoccupied, it remains on its current position. If it remains on food, the motion update is computed as if the agent is not at the food border with the exception that agents can only choose lattice sites with food to move to.

The motion updates of N2 agents are slightly different. If not located on food, an agent performs a random step to a lattice site in its remote neighbourhood. On food, it slows down and moves to a lattice site in its direct neighbourhood. If there is food on every lattice site in its direct neighbourhood, it performs a random step. If the agent is located at the food border, however, it decides whether to remain or leave the food according to ρ. In the former, it randomly chooses one of the lattice sites with food in the direct neighbourhood; in the latter, it chooses a lattice site without food in the direct neighbourhood. Only if no available lattice site fulfils these criteria, the agent remains at its position.

#### *Calculating food-leaving probability from experimental data*

Food-leaving events are identified from single-spot experiments for both strains, and are very rare on the thin food lawns that we use, compared to what others have reported under different experimental conditions (1, 2). Because food-leaving frequency increases as food becomes increasingly depleted (1), and because food is depleted at different rates between the two strains,(3) we sampled food-leaving events from the recording durations that correspond to 40-60% into the total food depletion duration. Food-leaving probability is obtained by dividing the total number of food leaving events during the sample duration by the length of the duration and then by the median number of worms on the food spot over the duration.

### Supplementary References

1. Milward K, Busch KE, Murphy RJ, de Bono M, Olofsson B. Neuronal and molecular substrates for optimal foraging in Caenorhabditis elegans. Proceedings of the National Academy of Sciences of the United States of America. 2011;108(51):20672-7.
2. Shtonda BB, Avery L. Dietary choice behavior in Caenorhabditis elegans. J Exp Biol. 2006;209(Pt 1):89-102.
3. Ding SS, Romenskyy M, Sarkisyan KS, Brown AEX. Measuring Caenorhabditis elegans Spatial Foraging and Food Intake Using Bioluminescent Bacteria. Genetics. 2020.
